# How phenotypic convergence arises in experimental evolution

**DOI:** 10.1101/579524

**Authors:** Pedro Simões, Inês Fragata, Josiane Santos, Marta A. Santos, Mauro Santos, Michael R. Rose, Margarida Matos

**Affiliations:** cE3c – Centre for Ecology, Evolution and Environmental Changes, Faculdade de Ciências, Universidade de Lisboa, Campo Grande, 1749-016 Lisboa, Portugal; Department of Ecology and Evolutionary Biology, University of California, Irvine, CA, USA; Departament de Genètica i de Microbiologia, Grup de Genòmica, Bioinformàtica i Biologia Evolutiva (GGBE), Universitat Autonòma de Barcelona, Spain

**Keywords:** Convergent Evolution, Experimental Evolution, *Drosophila subobscura*, Life-History traits, Long-term Evolution, Adaptation

## Abstract

Evolutionary convergence is a core issue in the study of adaptive evolution, as well as a highly debated topic at present. Few studies have analyzed this issue using a “real-time” or evolutionary trajectory approach. Do populations that are initially differentiated converge to a similar adaptive state when experiencing a common novel environment? *Drosophila subobscura* populations founded from different locations and years showed initial differences and variation in evolutionary rates in several traits during short-term (∼20 generations) laboratory adaptation. Here we extend that analysis to 40 more generations to analyze (1) how differences in evolutionary dynamics between populations change between shorter and longer time spans, and (2) whether evolutionary convergence occurs after sixty generations of evolution in a common environment. We found substantial variation in longer-term evolutionary trajectories and differences between short and longer-term evolutionary dynamics. Though we observed pervasive patterns of convergence towards the character values of long-established populations, populations still remain differentiated for several traits at the final generations analyzed. This pattern might involve transient divergence, as we report in some cases, indicating that more generations should lead to final convergence. These findings highlight the importance of longer-term studies for understanding convergent evolution.

## Introduction

Understanding how populations adapt to environmental challenges is becoming increasingly important in both evolutionary biology and conservation (Botero et al. 2015; Franks and Hoffmann 2012). However, we are still unsure how predictable adaptation to novel environments is (Lachapelle et al. 2015; Lässig et al. 2017; Lenski et al. 2015; Orgogozo 2015; Wiser et al. 2013). Unpredictability in evolution can be caused by different genetic backgrounds due to prior evolutionary history (see Barton and Keightley 2002; Barrett and Schluter 2008; Hansen 2013), and stochastic events such as founder events, genetic drift, bottlenecks, etc. (see Lenormand et al. 2009). Furthermore, interactions between selection and genetic drift may also increase variation in evolutionary responses (e.g. Cohan 1984; Cohan and Hoffmann 1986; Santos et al. 2012).

An important question when different populations adapt to new environmental challenges is whether they will diverge or converge through time. Convergent evolution is expected to arise through the action of natural selection, erasing differences between populations (Endler 1986; Losos 2011; Stern 2013). Alternatively, differentiated populations could conceivably evolve increased differentiation when placed under similar selective regimes (Wright 1931; Cohan 1984; Whitlock et al. 1995). Discovering the constraints that produce either evolutionary convergence or evolutionary divergence is fundamental to ultimately understanding the foundations of adaptive evolution.

Experimental evolution is a powerful tool with which to address this problem, especially by studying the real-time evolutionary trajectories of different populations subjected to the same selective challenge. Several studies have observed convergent evolutionary responses in a new common environment (e.g. Travisano et al. 1995; Teotónio and Rose 2000; 2002; Joshi et al. 2003; Simões et al. 2007, 2008; Teotónio et al. 2009; Santos et al. 2012; Fragata et al. 2014; Burke et al. 2016; Rebolleda-Gómez and Travisano 2019). Nevertheless, divergent evolutionary responses have also been observed (e.g. Cohan 1984; Cohan and Hoffmann 1986; Melnyk and Kassen 2011). Furthermore, several studies support the notion that the impact of evolutionary contingencies varies between traits closely or loosely related to fitness (Travisano et al. 1995; Teotónio et al. 2002; Joshi et al. 2003; Simões et al. 2008, 2017). It is thus clear from experimental evidence that evolutionary contingencies have a role in shaping evolutionary responses.

An important question, seldom addressed in the literature (but see Burke et al. 2016), is the effect of initial differentiation between populations on their long-term evolution. In particular, it is expected that different initial genetic backgrounds will have a higher impact during short-term evolution in a constant environment (Joshi et al. 2003; Fragata et al. 2014; Burke et al. 2016). On the other hand, at longer evolutionary scales, the cumulative effects of genetic drift and other stochastic events acting on the evolving populations will likely have a higher impact on the evolutionary trajectories observed (e.g. see Brito et al. 2005; Lenormand et al. 2009). Furthermore, different levels of standing genetic variation and/or epistatic interactions can have an important impact on long-term evolution (Barrett and Schluter 2008; Goodnight 2015; Paixão and Barton 2016; see empirical examples in Barton and Keightley 2002; Hansen 2013; Wiser et al. 2013; Good and Desai 2015). This might produce differences between populations, even in populations subject to similar selective pressures, possibly through different timings in the deceleration of the evolutionary response over time, for example (Teotónio and Rose 2000; Gilligan and Frankham 2003; Simões et al. 2007; Khan et al. 2011; Schoustra et al. 2012).

Long-term evolutionary dynamics have been mostly studied in microbial experimental evolution systems rather than in sexual organisms, due to the shorter generation time of the former. In the *E. coli* long-term evolution experiment performed in Lenski’s lab, recent evidence indicates a deceleration of the evolutionary rate over 50000 generations (Wiser et al. 2013; Lenski et al. 2015). Furthermore, and perhaps surprisingly, heterogeneity in evolutionary trajectories is still present after so many generations, in part due to differences in mutation rates (Lenski et al. 2015). Several studies with sexual organisms, though involving fewer generations, have also observed the slowing down of evolutionary responses to newly imposed selection regimes (e.g. Gilligan and Frankham 2003; Rose et al. 2004; Simões et al. 2007, see below). The expectation of a deceleration of laboratory evolutionary trajectories in sexual organisms is sometimes justified in terms of temporal exhaustion of additive genetic variance, although genomic scans in experimentally evolved *Drosophila* populations have found only limited evidence of fixed alleles following selection (Burke et al. 2010; Burke and Long 2012; Orozco-Terwengel et al. 2012; Long et al. 2015; Phillips et al. 2016; Seabra et al. 2018). In a previous study by our team, we found evidence for a deceleration in the evolutionary trajectory of fecundity in populations of *Drosophila subobscura* evolving for more than 80 generations in the lab environment (Simões et al. 2007). Teotónio and Rose (2000) also found this pattern of response in several *D. melanogaster* lines undergoing reverse selection in their ancestral environment. Gilligan and Frankham (2003) also reported a slowing down of the rate of adaptation to captivity after 87 generations in the lab by comparing *Drosophila* populations in different stages of adaptation. However, this pattern is not universal, as other experimental studies have not found such deceleration of the evolutionary response, even after a higher number of generations. One example of this is the work of Chippindale et al. (1997), who imposed selection for accelerated development time in *D. melanogaster*. Nevertheless, studies of long-term experimental evolution in sexual species are scarce and have not specifically addressed the variation in evolutionary dynamics that might occur during evolution over the short term, relative to longer evolutionary time periods.

We have previously shown variation in the evolutionary response of several populations of *D. subobscura* during the first 20 generations of evolution in a new environment, the lab (Simões et al. 2008). These populations were founded from different nearby locations over several years. We observed higher variation in the evolutionary response for female starvation resistance, a trait likely more loosely related to fitness in our experimental setting. By contrast, patterns for fecundity traits, which are expected to be closer to fitness, were more repeatable. Importantly, the different starvation resistance patterns led in fact to convergence between populations. In this study we extend the earlier analysis to cover around forty additional generations. We address the following questions: (1) How much do evolutionary rates vary between short-term and longer-term evolution? (2) Do differences in evolutionary dynamics between populations change in the transition from earlier to later generations? (3) Is convergence observed after sixty generations of evolution in a common environment?

We expect that, during short-term evolution, variation in the initial genetic backgrounds will lead to disparate rates of adaptation to the new environment. Over the longer term, as the evolutionary response decelerates, differences between populations of contrasting initial genetic composition are likely to be reduced relative to those observed during short-term evolution, particularly if populations are evolving towards the same phenotypic optimum.

## Materials and Methods

### Founding and Maintenance of the Laboratory populations

Five sets of wild-caught samples of *Drosophila subobscura* were analyzed in this study. These populations were founded in 1998 (NW populations; see Matos et al. 2002), 2001 (AR and TW populations; see (Simões et al. 2007), and 2005 (FWA and NARA; see (Simões et al. 2008). NW, TW and FWA populations were collected from a pinewood near Sintra (Portugal), whereas AR and NARA populations were collected from a pinewood in Arrábida (also from Portugal, some 50 Km from Sintra, on the other margin of the Tagus river; see Simões et al. 2007, 2008). All populations were three-fold replicated two generations after founding (e.g., FWA_1-3_ designating the three populations of FWA). A set of long-established laboratory populations (called “NB”, founded in 1990 from Sintra) was used as a control for all the experimental populations. NB populations were at their 90^th^, 136^th^ and 181^st^ laboratory generations at the time of foundation of the 1998, 2001 and 2005 collections, respectively.

All populations were maintained under the same laboratory environment with discrete generations of 28 days, reproduction close to peak fecundity, controlled temperature of 18°C, with a 12-h L: 12-h D photoperiod. Flies were kept in vials, with controlled densities for both adult (around 50 individuals per vial) and larval stages (around 80 per vial). At each generation, emergences from the several vials of each replicate population were randomized using CO_2_ anesthesia. Census population sizes ranged between 600 and 1200 adults. To study the evolutionary trajectories during laboratory adaptation, all experimental populations and the controls were periodically assayed for several phenotypic traits (see below).

### Phenotypic Assays and Generations analyzed

For the phenotypic assays, mated pairs of flies were transferred daily to fresh medium and the number of eggs laid per female was counted during the first 12 days since emergence. After the fecundity assay, each pair of flies was transferred to a vial containing plain agar medium to measure starvation resistance (with deaths checked every 6 h). Five characters were analyzed: age of first reproduction (number of days between emergence and the day of first egg laying), early fecundity (total number of eggs laid during the first week), peak fecundity (total number of eggs laid between days 8 and 12), and female and male starvation resistance. Sample sizes ranged between 14 and 24 pairs per replicate population and assay. All assays involved synchronous analyses with NB populations.

Periodical phenotypic assays were performed starting at generation 3 or 4 up to generation 58-60. All generations assayed for the several populations are presented in Table S1. We analyze here both short-term - ∼20 generations - and a longer-term period-between ∼20 and ∼60 generations, here designated “long-term” - of laboratory evolution of these populations. We also analyzed the entire evolutionary trajectory, spanning the complete data set. The short-term data was studied in Simões et al. (2008) for a larger number of populations, the five sets of populations referred to above and an extra set of populations in each of the 2005 locations (details in Simões et al. 2008). Moreover, for NW there were five replicate populations with data on short term, but here we only analyze three replicate populations, for both short and long-term, as only these have data for more advanced generations. Finally, we expand our analyses to include male starvation resistance data, which was not analyzed in Simões et al. (2008).

In order to calculate the initial or final state for each replicate population, we calculated the mean value of the 2 (or 3) first (or last) generations by choosing 15 random individual data points (with replacement) of each generation involved. The initial generations used were the following: 4, 6 and 7 for AR and TW; 4 and 8 for NW; 3, 6 and 10 for NARA and FWA. The final generations analyzed were: 48, 55 and 60 for AR and TW; 52, 53 and 58 for NW; 49 and 58 for NARA and FWA.

### Statistical Methods

To estimate the evolutionary trajectories for each population, in each assayed generation, we used the differences between individual data and the mean of the same-numbered NB replicate population (assayed synchronously with experimental populations; e.g. AR1-average NB1), see (Simões et al. 2008). This was done to remove the effect of possible temporal changes not related to laboratory adaptation such as trends due to environmental variation or to inadvertent evolutionary changes not intended in the study (e.g. due to slight changes of conditions in lab). This procedure also minimizes the effects of environmental heterogeneity between non-synchronous assays (see also Matos et al. 2002; Simões et al. 2007, 2008). Temporal performance of the control populations was generally quite stable across traits, allowing us to rule out undesirable sources of variation such as those due to further laboratory adaptation or inbreeding (see Fig S1).

Linear and linear-log models were tested for both periods separately and over the whole evolutionary trajectory of the populations (around 60 generations). Models were chosen according to their fit to the data based on R^2^ values (see Table S2). For the separate analyses of short-term and long-term periods, we chose the linear over the linear-log model as a compromise across populations and periods, since the same model had to be applied to allow for direct comparisons between periods (e.g. for the tests in Table 1 and 2). For the analysis of overall trajectories, the linear-log model was chosen, as it generally presented a better fit than the linear model (see Table S2).

**Table 1.**
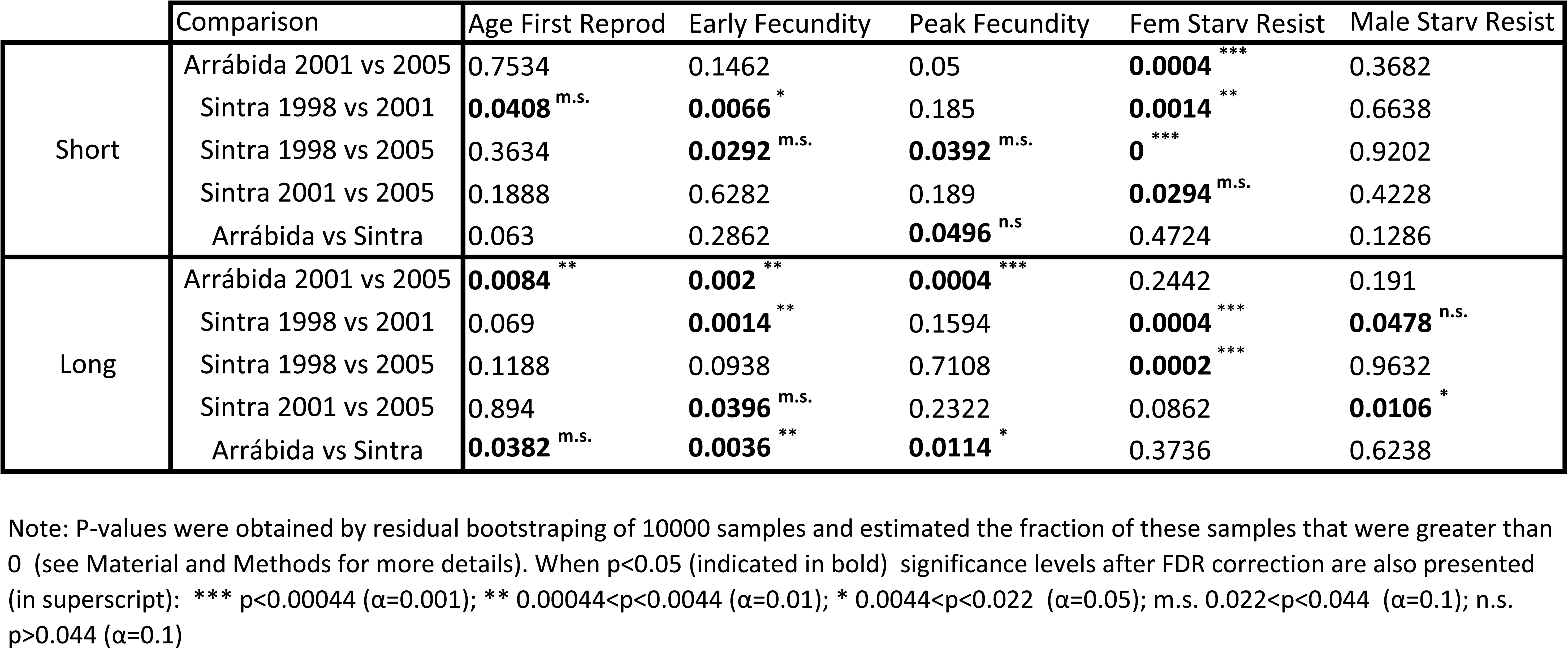
Comparison of evolutionary rates between different years or locations for short or longer periods.

|  | Comparison | Age First Reprod | Early Fecundity | Peak Fecundity | Fem Starv Resist | Male Starv Resist |
| --- | --- | --- | --- | --- | --- | --- |
| Short | Arrábida 2001 vs 2005 | 0.7534 | 0.1462 | 0.05 | **0.0004**<sup>***</sup> | 0.3682 |
|  | Sintra 1998 vs 2001 | **0.0408**<sup>m.s.</sup> | **0.0066**<sup>*</sup> | 0.185 | **0.0014**<sup>**</sup> | 0.6638 |
|  | Sintra 1998 vs 2005 | 0.3634 | **0.0292**<sup>m.s.</sup> | **0.0392**<sup>m.s.</sup> | **0**<sup>***</sup> | 0.9202 |
|  | Sintra 2001 vs 2005 | 0.1888 | 0.6282 | 0.189 | **0.0294**<sup>m.s.</sup> | 0.4228 |
|  | Arrábida vs Sintra | 0.063 | 0.2862 | **0.0496**<sup>n.s.</sup> | 0.4724 | 0.1286 |
| Long | Arrábida 2001 vs 2005 | **0.0084**<sup>**</sup> | **0.002**<sup>**</sup> | **0.0004**<sup>***</sup> | 0.2442 | 0.191 |
|  | Sintra 1998 vs 2001 | 0.069 | **0.0014**<sup>**</sup> | 0.1594 | **0.0004**<sup>***</sup> | **0.0478**<sup>n.s.</sup> |
|  | Sintra 1998 vs 2005 | 0.1188 | 0.0938 | 0.7108 | **0.0002**<sup>***</sup> | 0.9632 |
|  | Sintra 2001 vs 2005 | 0.894 | **0.0396**<sup>m.s.</sup> | 0.2322 | 0.0862 | **0.0106**<sup>*</sup> |
|  | Arrábida vs Sintra | **0.0382**<sup>m.s.</sup> | **0.0036**<sup>**</sup> | **0.0114**<sup>*</sup> | 0.3736 | 0.6238 |

Note: P-values were obtained by residual bootstrapping of 10000 samples and estimated the fraction of these samples that were greater than 0 (see Material and Methods for more details). When $p < 0.05$ (indicated in bold) significance levels after FDR correction are also presented (in superscript): \*\*\* $p < 0.00044$ ( $\alpha = 0.001$ ); \*\* $0.00044 < p < 0.0044$ ( $\alpha = 0.01$ ); \* $0.0044 < p < 0.022$ ( $\alpha = 0.05$ ); m.s. $0.022 < p < 0.044$ ( $\alpha = 0.1$ ); n.s. $p > 0.044$ ( $\alpha = 0.1$ )

**Table 2.** Comparison of short and long term evolutionary rates between years and locations.

| Comparison | Age First Reprod | Early Fecundity | Peak Fecundity | Fem Starv Resist | Male Starv Resist |
| --- | --- | --- | --- | --- | --- |
| Arrábida 2001 vs 2005 | 0.2376 | **0.0042**<sup>**</sup> | 0.7132 | **0**<sup>***</sup> | 0.1444 |
| Sintra 1998 vs 2001 | 0.1402 | **0**<sup>***</sup> | 0.069 | **0**<sup>***</sup> | 0.2288 |
| Sintra 1998 vs 2005 | 0.6716 | **0.006**<sup>*</sup> | **0.048**<sup>n.s.</sup> | **0**<sup>***</sup> | 0.9166 |
| Sintra 2001 vs 2005 | 0.238 | 0.1714 | 0.5928 | **0.0078**<sup>*</sup> | 0.0702 |
| Arrábida vs Sintra | 0.2628 | 0.6254 | 0.7414 | 0.2986 | 0.1144 |

Note: P-values were obtained by residual bootstrapping of 10000 samples and estimated the fraction of these samples that were greater than 0 (see Material and Methods for more details). Significant results are indicated in bold. When $p < 0.05$ (indicated in bold) significance levels after FDR correction are also presented (in superscript): \*\*\* $p < 0.00044$ ( $\alpha = 0.001$ ); \*\* $0.00044 < p < 0.0044$ ( $\alpha = 0.01$ ); \* $0.0044 < p < 0.022$ ( $\alpha = 0.05$ ); m.s. $0.022 < p < 0.044$ ( $\alpha = 0.1$ ); n.s. $p > 0.044$ ( $\alpha = 0.1$ )

### Bootstrap Techniques

Variation in the slope of evolutionary response between sets of populations and periods was studied using bootstrap techniques as in Simões et al. (2008). Briefly, for each replicate population we estimated the intercept 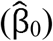, evolutionary slope 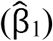 and the residuals of each point (ε) using a simple linear regression. In each iteration of the bootstrap, a new vector of phenotypic data was created by resampling the residuals, with replacement (ε*) and employing the following formula to calculate a new phenotypic value for each data point used:

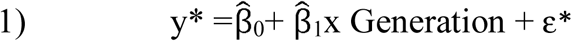

After this, a new slope (β_1_*) and intercept (β_0_*) were estimated through a linear regression. For the linear-log model the same analysis was applied using the natural logarithm of the generation. A total of 10000 slopes were generated for each replicate population. All analyses testing differences between slopes were done using these values.

To compare two sets of populations from the same location in different years, we calculated the mean of each set involved in the comparison (by randomly sampling one slope from each replicate population) and the difference between them (e.g. comparison Arrábida 2001 vs. Arrábida 2005: ((AR_1_β_1_* + AR_2_β_1_* + AR_3_β_1_*)/3) – ((NARA_1_β_1_*+NARA_2_β_1_*+NARA_3_β_1_*)/3). This process was repeated 10000 times. Statistical significance was assessed by estimating the fraction of these 10000 differences that were greater than zero. Two times this fraction or 1 minus two times this fraction (whichever is less) corresponds to the *P*-value. To compare differences between the two locations we used all 2001 and 2005 replicate populations from each location. NW data was not included as there were no corresponding populations founded in Arrábida in 1998. We calculated the location means using data of six replicate populations (e.g. FWA_1-3_ and TW_1-3_ for the Sintra slopes), again using random samples of slopes from each replicate population, as above. Differences between the short and long-term evolutionary response for each set of populations were also assessed (e.g. comparison of TW_1-3_ short-term slopes vs TW_1-3_ long-term slopes). We further analyzed whether differences between periods varied between populations founded from distinct years or locations. These comparisons followed the same rationale as above (e.g. comparison short vs long-term for Arrábida 2001 vs Arrábida 2005: ((AR_1_β_1_*_S_+ AR_2_β_1_*_S_+AR_3_β_1_*_S_)/3 – (NARA_1_β_1_*_S_ + NARA_2_β_1_*_S_ + NARA_2_β_1_*_S_)/3)-((AR_1_β_1_*_L_+ AR_2_β_1_*_L_+AR_3_β_1_*_L_)/3 – (NARA_1_β_1_*_L_ + NARA_2_β_1_*_L_ + NARA_2_β_1_*_L_)/3). This analysis was performed with 10000 random samples and tested as described above.

To test whether populations differed in the initial or final performance, 10000 comparisons between years and locations were assessed using the same rationale as above.

When testing for differences between populations statistical significance is presented both with and without False Discovery Rate (FDR) correction for five tests (theorem 1.3 Benjamini and Yekutieli 2001). Marginally significant results after FDR correction will also be considered when the general reading justifies, i.e. if there are consistent patterns across populations. This is a compromise, as being too conservative also has drawbacks, given that the focus of this study is not on single tests but rather to analyze patterns across comparisons.

All analyses were performed using R version 3.3.1 (R Core Team 2018), package reshape2 (Wickham 2007) and visualization was done using ggplot2 package (Wickham 2009).

## Results

### Initial differences between populations

The experimental populations were clearly differentiated from the control populations in the initial performance of fecundity traits, though less so for starvation resistance (Fig 1 and 2). NW populations performed significantly better than the other Sintra populations, both in age of first reproduction and early fecundity, whereas they performed worse for male starvation resistance (see Table S3 and Figs 1 and 2). Most populations from different years showed significant differences in the initial performance for peak fecundity and female starvation resistance. On the other hand, no significant differences were found between locations for any trait (see Table S3).

**Figure 1.**
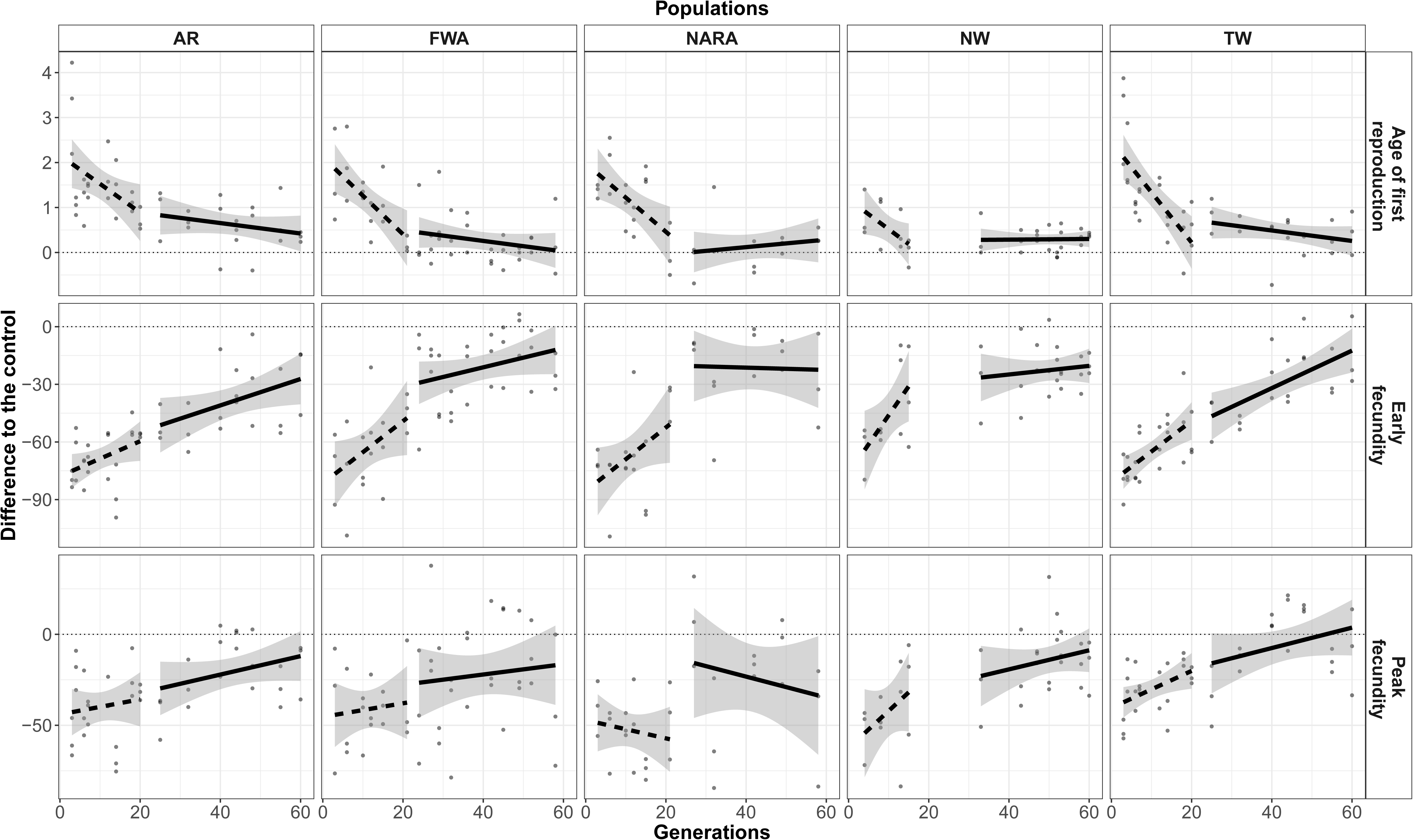
Short and long-term evolutionary trajectories for fecundity related traits for the 5 sets of populations studied. Age of first reproduction (number of days), Early fecundity (number of eggs), Peak fecundity (number of eggs) are represented. Points represent mean values for each replicate at each generation. Dashed lines indicate short term period and full line indicates long-term period. Shaded area represents 95% confidence intervals estimated from the regression, using mean replicate population values.

**Figure 2.**
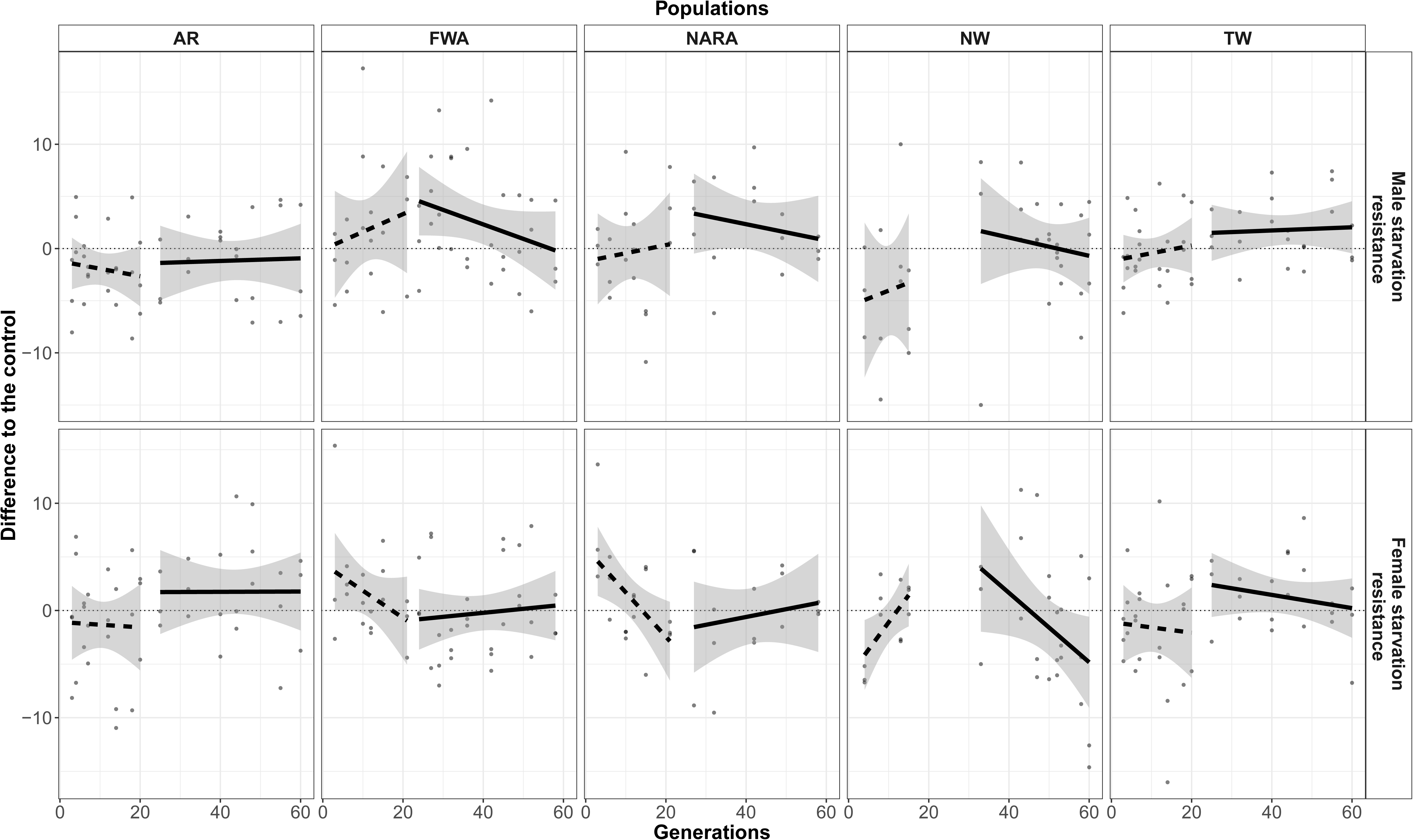
Short and long-term evolutionary trajectories for female and male starvation resistance for the 5 sets of populations studied. Male starvation resistance (in hours) and Female starvation resistance (in hours) are represented. Points represent mean value for each replicate at each generation. Dashed lines indicate short term period and full line indicates long-term period. Shaded area represents 95% confidence intervals estimated from the regression, using mean replicate population values.

### Short-term Evolutionary Dynamics

In general, fecundity-related traits, particularly early fecundity, show a clear evolutionary increase in performance during short-term evolution across populations, with a tendency to converge to control values, although at different rates (see Fig 1 and below; see also Fig S2, for data on the mean and variation of slopes of replicate populations). In contrast, patterns for starvation resistance are less consistent. In fact, male starvation resistance does not show a noticeable evolutionary response, although there is a suggestion of increased starvation across generations for all sets of populations except AR (see Fig 2 and Fig S2). The evolutionary response of female starvation resistance varies greatly among sets of populations with patterns of stasis (AR and TW), decreased (FWA and NARA) and increased performance (for NW). In spite of these differences, the patterns are again of convergence to control values (see Fig 2, S2 and below).

When comparing the evolutionary response among populations, we observe significant differences of slopes between years (see Table 1). This variation is particularly evident for female starvation resistance, in agreement with our previous analysis (Simões et al. 2008). On the other hand, for male starvation resistance no significant variation in the evolutionary response was found. Significant differences between locations were only observed for peak fecundity (see Table 1).

Interestingly, of the eight comparisons showing significant (or at least marginally significant, after FDR correction) differences between populations in short-term dynamics across all assayed traits (Table 1), six of these showed also significant variation in initial performance (cf. Table 1 and Table S3). This concordance corresponded to a reduction of differences between populations through time for age of first reproduction and female starvation resistance. In contrast, for early fecundity, a higher initial performance of NW relative to TW or FWA was followed by faster improvement through time increasing the initial differences, leading at least to transient divergence between populations (Table 1 and Table S3, Fig 1 and 2).

### Long-term Evolutionary Dynamics

In each set of populations there was a clear variation of evolutionary rates (slopes) between the short-term and the long-term period for age of first reproduction, early fecundity and female starvation resistance (Table S4). This corresponded to a general slowing down of the evolutionary response as populations tended to converge to the control values (see Figs 1, 2 and S2; Fig S3 shows the same pattern in the evolutionary trajectories using all generations).

Differences in evolutionary dynamics between sets of populations were more evident in the long-term than in the short-term evolutionary response for several traits, particularly for early fecundity (see Table 1; Figs 1, 2 and S2). For this trait, a significant effect of location was due to a higher evolutionary rate in Sintra populations. Also, several comparisons showed significant effects of year, due in part to a lower slowing down of the response of the 2001 populations. On the other hand, differences between populations in the evolutionary response of female starvation resistance decreased in this period with only two significant effects in five comparisons– see Table 1. These significant effects involved comparisons with NW, which showed a clear drop in performance during this later period (see Fig 2).

When comparing the variation in evolutionary rates of the different sets of populations between the two periods (short vs. long-term evolution), early fecundity and female starvation resistance showed the greatest differences between populations, due to the above mentioned differential slowing down of response for early fecundity during long-term evolution and to the reported high variation in evolutionary rates seen in the short term evolution of female starvation resistance (Figs 1 and 2, Table 2). Importantly, for early fecundity, populations with higher short-term evolutionary rates (NW and the two 2005 populations) were also those with a stronger slowing down in the long-term period (Fig 1), which is expected under convergent evolution (see below).

### Final differences between populations

In more advanced generations, there was a loss of the initial differences between populations for several comparisons, as expected if full convergence occurs (see Table S3). This was observed between NW and TW for all fecundity traits, between NW and FWA for age of first reproduction, and between the 2001 and 2005 populations for female starvation resistance (Table S3 and Figs. 1 and 2). Nevertheless, significant differences in final performance were also found for several comparisons (see Table S3). In some cases, differentiation was also present at the start. Three temporal patterns were observed taking into account initial, intermediate, and final values (see Table S5): 1-continuous reduction of differences (NW versus TW for male starvation resistance); 2-increased differences through time (TW versus FWA for peak fecundity); 3-differentiation at the initial and final generations but with intermediate loss of differentiation (NW versus FWA for early fecundity and female starvation resistance). Finally, in other comparisons there was a significant (or at least marginally significant after FDR correction) differentiation between populations at the later stage of adaptation, not present at the start (Table S3). In this case two temporal patterns were observed (see Fig 1 and Table 1, S3 and S5): 1-higher differences at the end than at intermediate or initial generations (Arrábida versus Sintra populations for early fecundity and NW versus FWA for peak fecundity); 2 - - higher differences at intermediate generations than at the end of the study due to a differential slowing down of the evolutionary rate (between the two sets of Arrábida populations for age of first reproduction and early fecundity; in both cases differences are marginally significant after FDR correction).

### Overall Evolutionary Dynamics

Evolutionary trajectories across the entire time span confirm a general deceleration of the evolutionary response through time, as populations evolved towards the control values (see Fig S3). This led to a generally better fit of the overall evolutionary trajectory to a linear-log model relative to a linear one, particularly for fecundity-related data (see Table S2). Differences between sets of populations in the overall evolutionary response were due to variable changes between short and long periods, leading to pervasive contrasts, particularly for early fecundity (see Table 1 and S6).

## Discussion

Evolutionary convergence is a core expectation for adaptive evolution in a similar environment (Losos 2011; Stern 2013). With a smooth fitness landscape, that lacks multiple peaks, populations will tend to evolve to the same outcome (Wright 1931). In such cases, the outcome of evolution will be predictable. The predictability of evolution is an issue of much interest at present (e.g. de Visser and Krug 2014; Orgogozo 2015). Experimental evolution is a great tool for testing whether adaptive evolution involves smooth or rugged landscapes, as it allows us to study the fate of populations initially differentiated when subject to similar selective pressures, especially whether they evolve towards similar or different fitness values (Fragata et al. 2018; Matos et al. 2015; Orgogozo 2015; Rebolleda-Gómez and Travisano 2019). Here we add to the previous Simões et al. (2008) study the analysis of c. 40 more generations of laboratory adaptation, in order to determine whether: 1) longer-term evolution leads to similar outcomes as short-term evolution; 2) populations will ultimately tend to converge or show more complex evolutionary patterns.

In this study we found a general pattern of convergent evolution, with clear changes in the evolutionary rates between the short-term (∼20 generations) and longer-term (∼60 generations) periods. We observed a slowing down of the evolutionary response through time for several traits as populations approached the evolutionary equilibria of long-established populations. Empirical evidence for deceleration of evolutionary rate has been observed in other experimental studies using both asexual (Wiser et al. 2013; Lenski et al. 2015) and sexual organisms (Gilligan and Frankham 2003; Simões et al. 2007).

We also observed that the differences between short-term and longer-term dynamics were trait and population specific. Whereas differences in the early-fecundity response between sets of populations increased from short- to long-term evolution, the inverse pattern was observed for female starvation resistance. The source of differences between populations also varied between traits. In the case of early fecundity, trajectory variation was due to a continuous increase in performance of the 2001 populations, even during long-term evolution, contrasting with the 1998 and 2005 populations, where quicker short-term evolution was followed by a slowing of the evolutionary response after generation 20. These differences are consistent with convergent evolution, as faster evolution in an earlier period is followed by a plateauing, while slower evolution corresponds to a steadier evolutionary rate throughout generations. Such contrasting evolutionary dynamics led to an interesting pattern: an intermediate phase of transient divergence was followed in the long-term by a partial convergence among evolving populations. In contrast, for female starvation resistance there were striking differences in the evolutionary trajectories during short-term evolution, with increase, decrease, or stasis contingent on the degree of initial differentiation from controls (see also Simões et al. 2008). For this trait, convergence was fast between all populations. These patterns were followed in general by a reduction of differences between evolutionary trajectories over the longer time period analyzed. The exception was the NW populations, which presented an initial positive trend, unique across populations (see also Matos et al. 2004), followed by a negative long-term trend. Nevertheless, despite the different underlying evolutionary dynamics, both early fecundity and female starvation resistance show a general pattern that suggests convergence in longer-term periods.

It is an inherent expectation of convergent evolution that there will be a negative association between initial state and subsequent evolutionary rates of populations adapting to a new environment (Simões et al. 2007). This expectation was confirmed for *D. subobscura* populations with clear initial historical differentiation, founded from contrasting latitudes of the European cline (Fragata et al. 2014). In that study fast convergence was observed after only 14 generations in a common environment. In our study, evidence of such an association was only found for age of first reproduction and female starvation resistance for the short-term dynamics. Even so, for female starvation resistance the overall trend was not of convergence in the case of NW populations (see above). The relative lack of such overall and rapid convergence in our study might be due to the smaller degree of initial differentiation of these populations, with greater sampling effects (Santos et al. 2012).

If full convergence occurs, an obvious corollary is that populations will not be differentiated as an outcome of evolution in a common environment. This expectation was not entirely met in our study, as several populations remained differentiated for some traits after sixty generations of evolution. In this context, several patterns emerged when comparing dynamics between different populations: (1) continuous reduction of differences indicating partial convergence (for male starvation resistance); (2) continuous divergence between populations (for early and peak fecundity); (3) transient divergence followed by partial convergence (for age of first reproduction and early fecundity) or (4) transient convergence followed by later divergence (for early fecundity and female starvation resistance). Teotónio and his collaborators (Teotónio et al. 2002; Teotónio and Rose 2000) performed a reverse evolution study during 50 generations involving many genetically differentiated *Drosophila melanogaster* populations. They found that populations converged to ancestral values, but this trend was not general as it varied with the previous history and the trait studied. They concluded that populations converged to similar fitness values to a larger extent than other characters did. In contrast, in our study we did not see any clear relation between the extent of convergence and how the traits analyzed were presumed to determine fitness. In fact, several populations remained differentiated for early fecundity, a trait that is under strong selection in our environment with clear and consistent improvement across many independent studies (Fragata et al. 2014; Matos et al. 2002; Matos et al. 2004; Simões et al. 2007; Simões et al. 2008). Given our interpretation of transient divergence and partial convergence in some of these populations, it is possible that the time span of the study was not sufficient to allow for full convergence in some cases, convergence that might ultimately occur over more generations of evolution.

We observed considerable differences between short-term and longer-term dynamics in all our populations, which raises questions about predicting long-term evolution from short-term evolution. This contrasts with the study of Burke et al. (2016), which suggests that short-term evolution is predictive of longer evolutionary time periods. In that study recently selected *D. melanogaster* populations converged to the trait values of other independently derived populations evolving in a similar selection regime for a longer time scale, regardless of the evolutionary history of the populations studied. However, different time scales were involved, as the shorter-term evolutionary responses of that study were sometimes more than 100 generations in duration, with long-term evolution approaching 1,000 generations. In general, the fact that our study showed such differentiated outcomes and complex evolutionary patterns highlights the importance of characterizing extended periods of experimental evolution and the possible pitfalls of predicting evolution from short-term adaptive patterns.

## Conclusions

We here showed that after 60 generations of evolution in a common environment, *Drosophila subobscura* populations remain differentiated for several traits. Noticeably, this was observed even for life-history traits that are clearly under selection in our lab. In this context, we found evidence for transient divergence, as a result of heterogeneity in evolutionary rates through time, occurring under a general scenario of convergence. Ultimately, we conclude that extrapolating from short-term evolutionary patterns to longer evolutionary periods might be risky, particularly if one is interested in predicting the outcomes of evolution.

## Acknowledgments

The authors thank Laurence D. Mueller for the first version of the script that was later adapted for the analyses of the present study. Two anonymous reviewers provided helpful comments on an earlier draft. This study was partially financed by “Fundação para a Ciência e a Tecnologia” (FCT) projects POCTI/BSE/33673/2000, POCI-PPCDT/BIA-BDE/55853/2004 and PTDC/BIA-BDE/65733/2006 and by the cE3c FCT Unit UID/BIA/00329/2013. PS and IF have postdoctoral FCT grants (SFRH/BPD/86186/2012 and within the project JPIAMR/0001/2016 respectively). MS is funded by grants CGL2017-89160P from Ministerio de Economía y Competitividad (Spain), and 2017SGR 1379 from Generalitat de Catalunya.

## Data Accessibility Statement

Raw data will be available at the Dryad Digital Repository upon acceptance.

