## Supplemental Figure 2 for "How phenotypic convergence arises in experimental evolution"

**Supplementary Figure 2 -** Variation in 1000 boostrapped slopes for each trait and replicate population analyzed in short (left panels) and long-term (right panels) evolution.

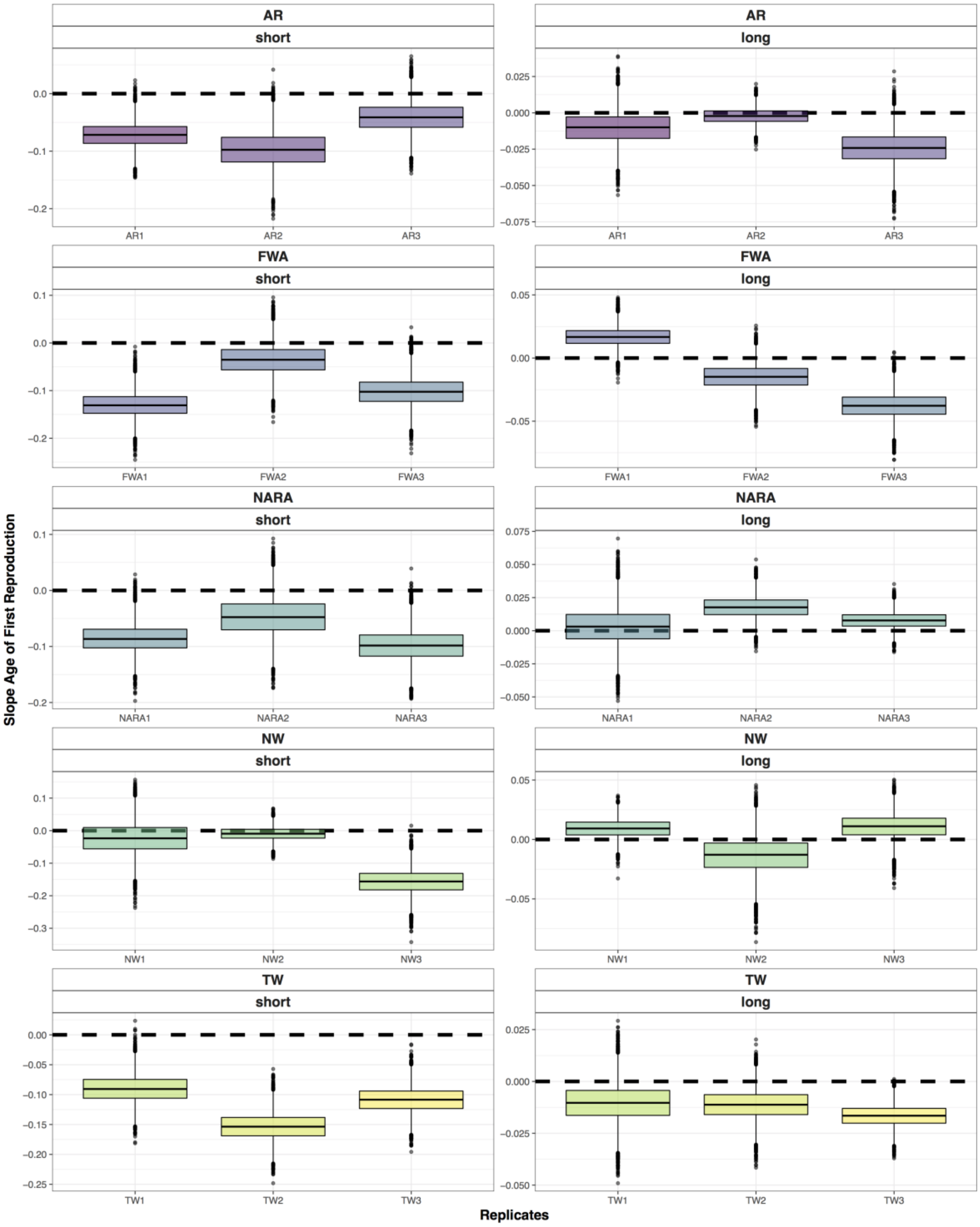


**A)**

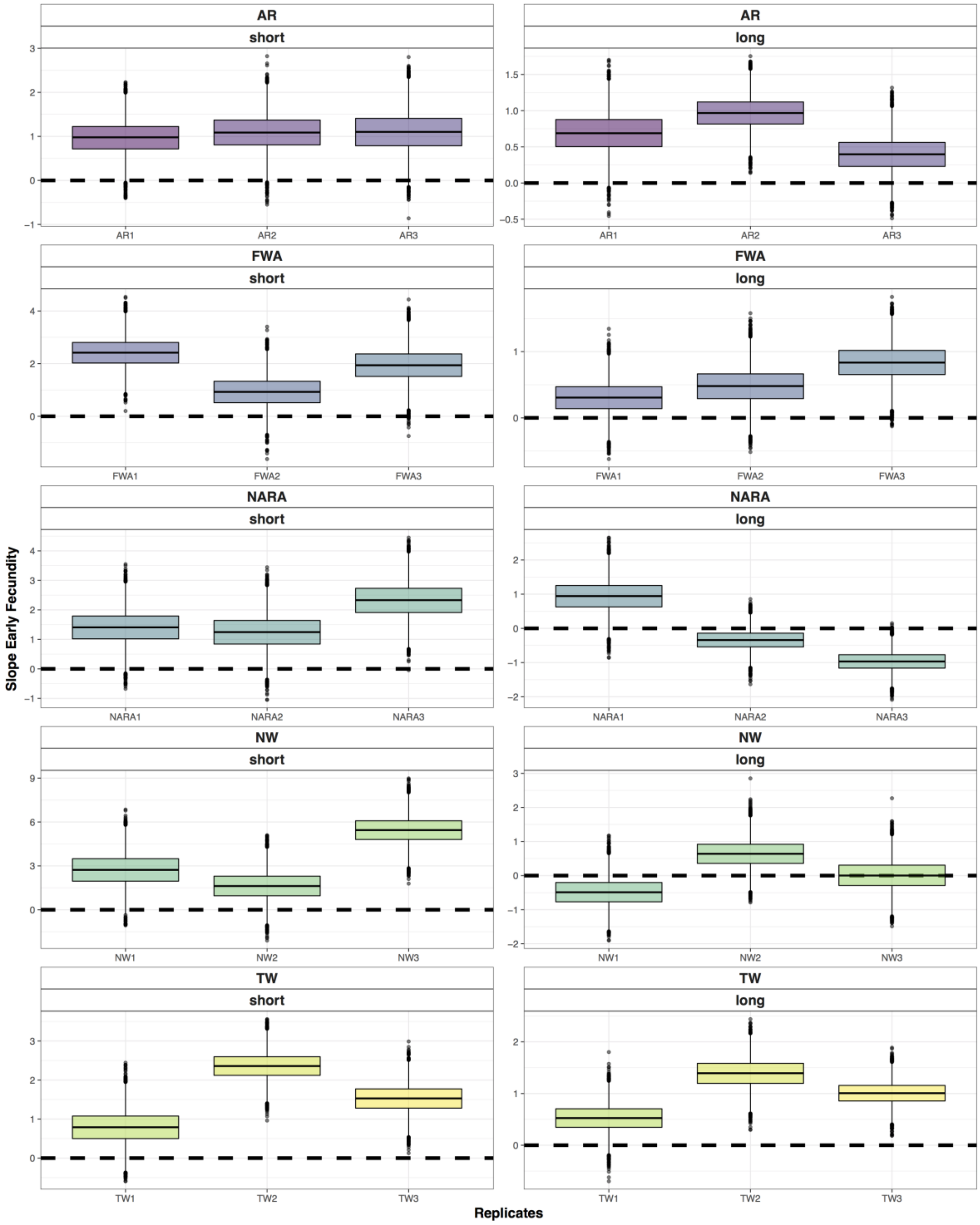
AR
short
long
AR1
AR2
AR3
FWA
FWA1
FWA2
FWA3
NARA
NARA1
NARA2
NARA3
NW
NW1
NW2
NW3
TW
TW1
TW2
TW3
Slope Early Fecundity
Replicates

**B)**

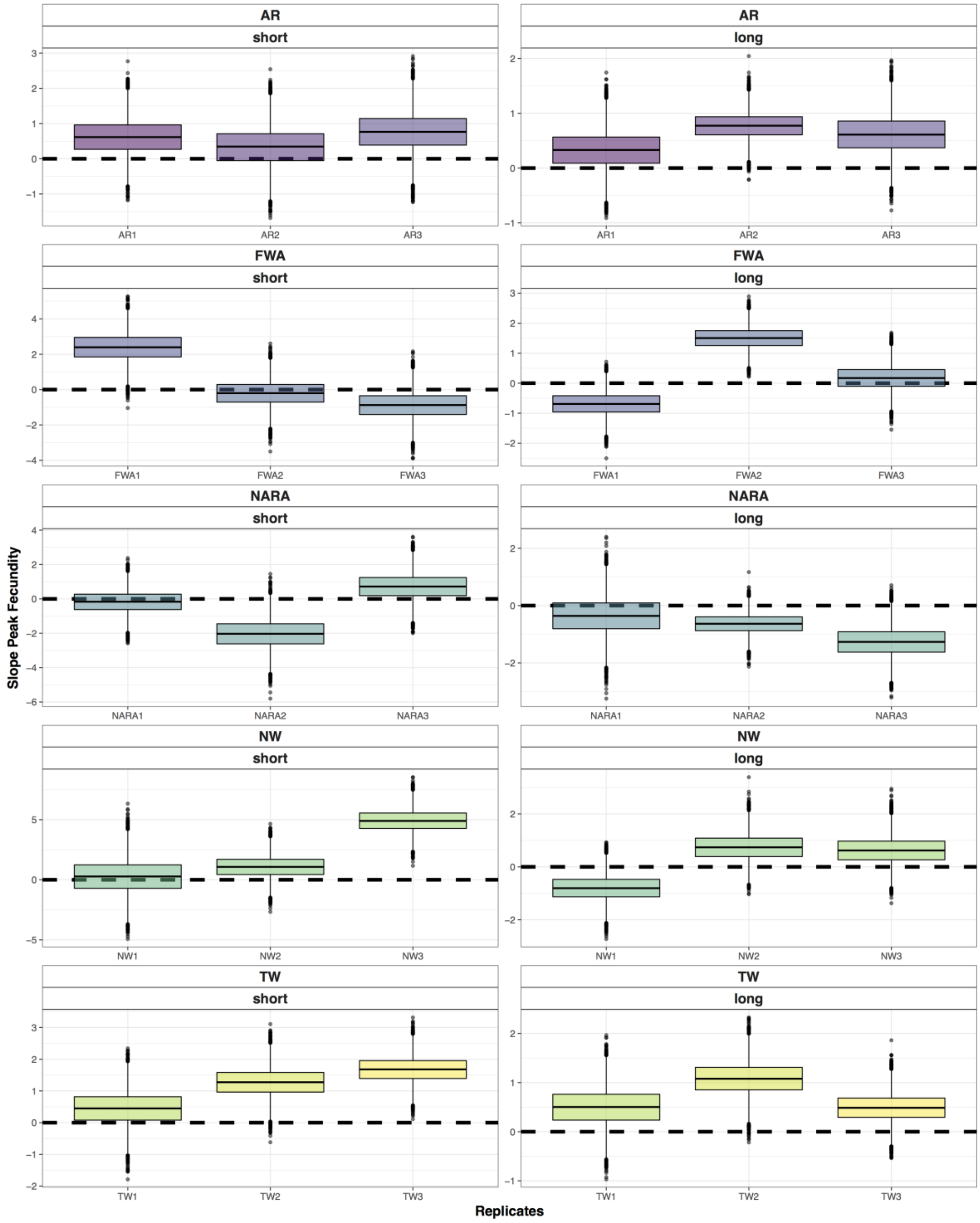
AR
short
AR
long
FWA
short
FWA
long
NARA
short
NARA
long
NW
short
NW
long
TW
short
TW
long
Slope Peak Fecundity
Replicates
AR1
AR2
AR3
FWA1
FWA2
FWA3
NARA1
NARA2
NARA3
NW1
NW2
NW3
TW1
TW2
TW3

C)

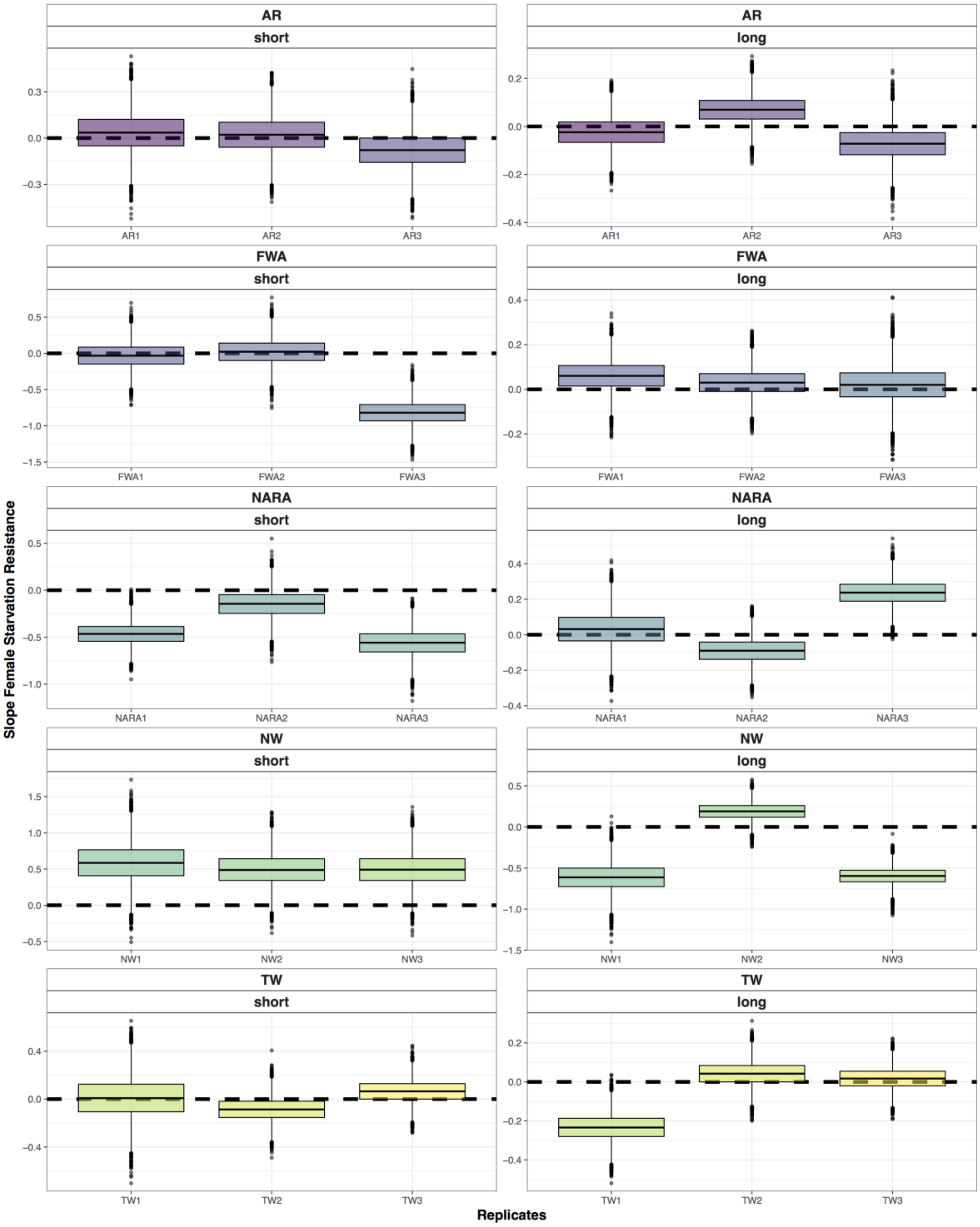
AR
short
AR
long
AR1
AR2
AR3
FWA
short
FWA
long
FWA1
FWA2
FWA3
NARA
short
NARA
long
NARA1
NARA2
NARA3
NW
short
NW
long
NW1
NW2
NW3
TW
short
TW
long
TW1
TW2
TW3
Slope Female Starvation Resistance
Replicates

**D)**

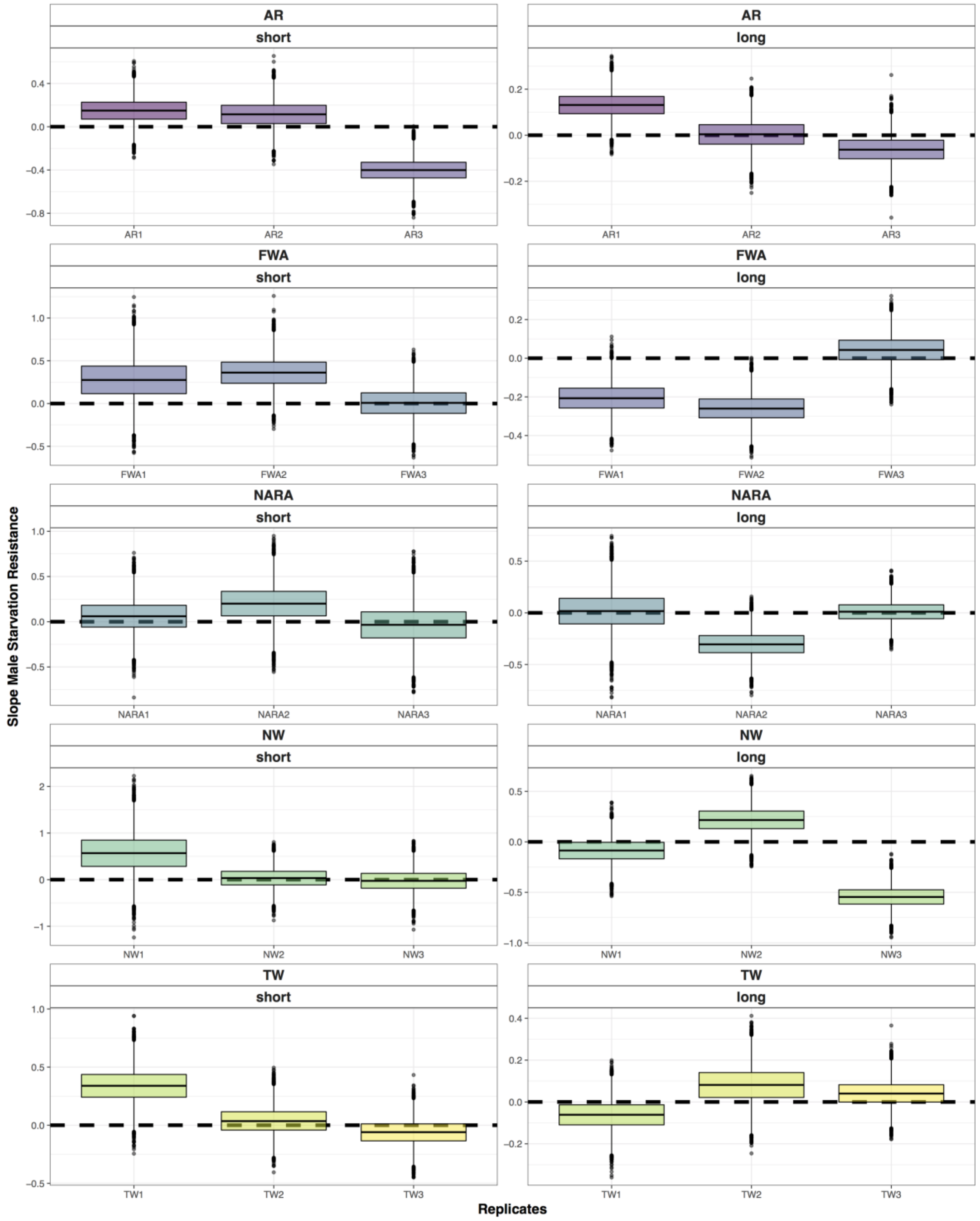


**E)**

Figure S2 - Variation in 1000 boostrapped slopes for Age of first reproduction (A), Early fecundity (B), Peak fecundity (C), Female starvation resistance (D) and Male starvation resistance (E) for replicates in short (left panels) and long-term (right panels) evolution. For each replicate population the box-plot represents the median and 95% confidence interval of the 1000 bootstrap slopes for short and long-term periods.
