## Supplementary figures and images for "How phenotypic convergence arises in experimental evolution"

### Supplemental Figure 1

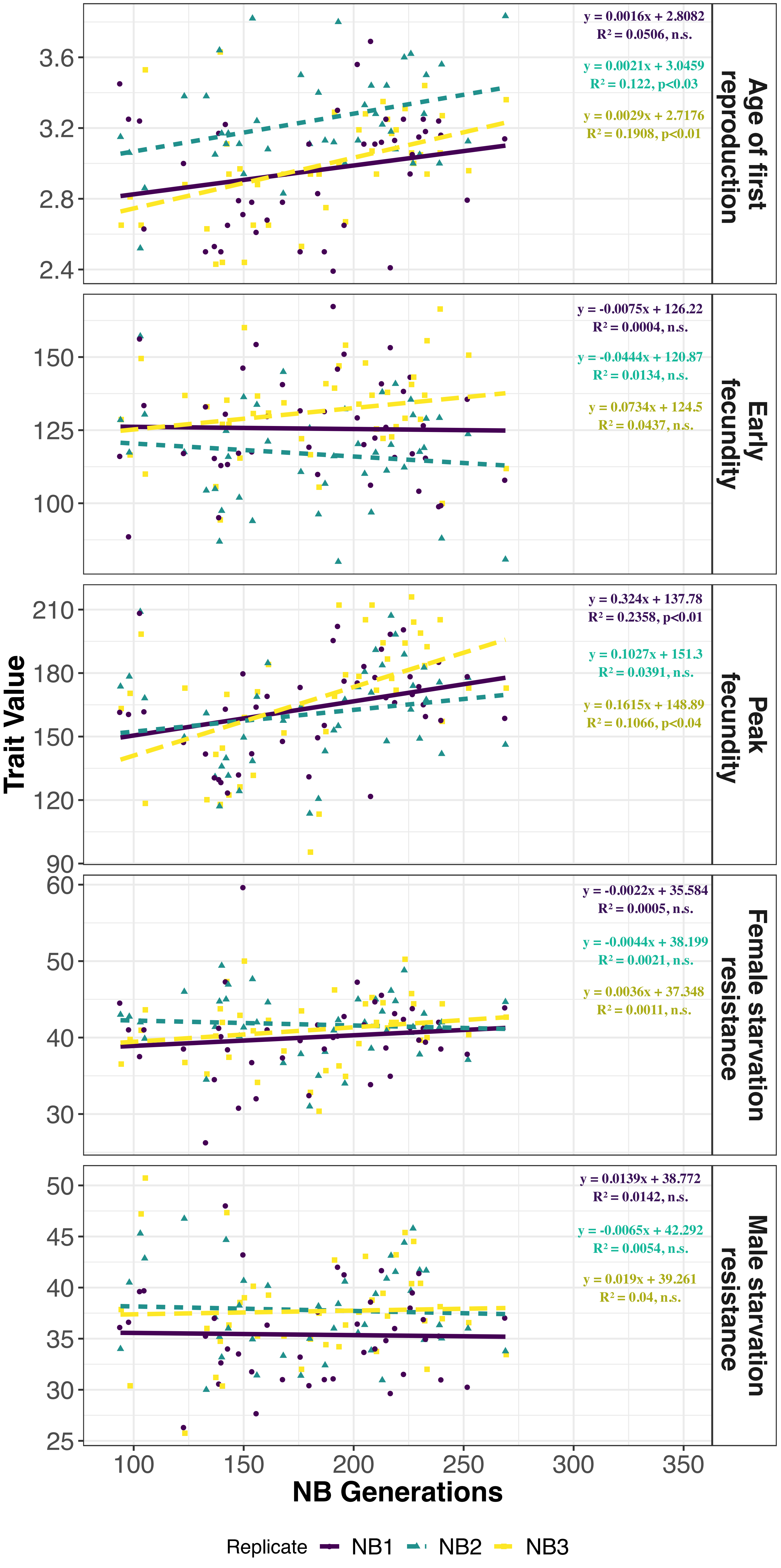

### Supplemental Figure 3

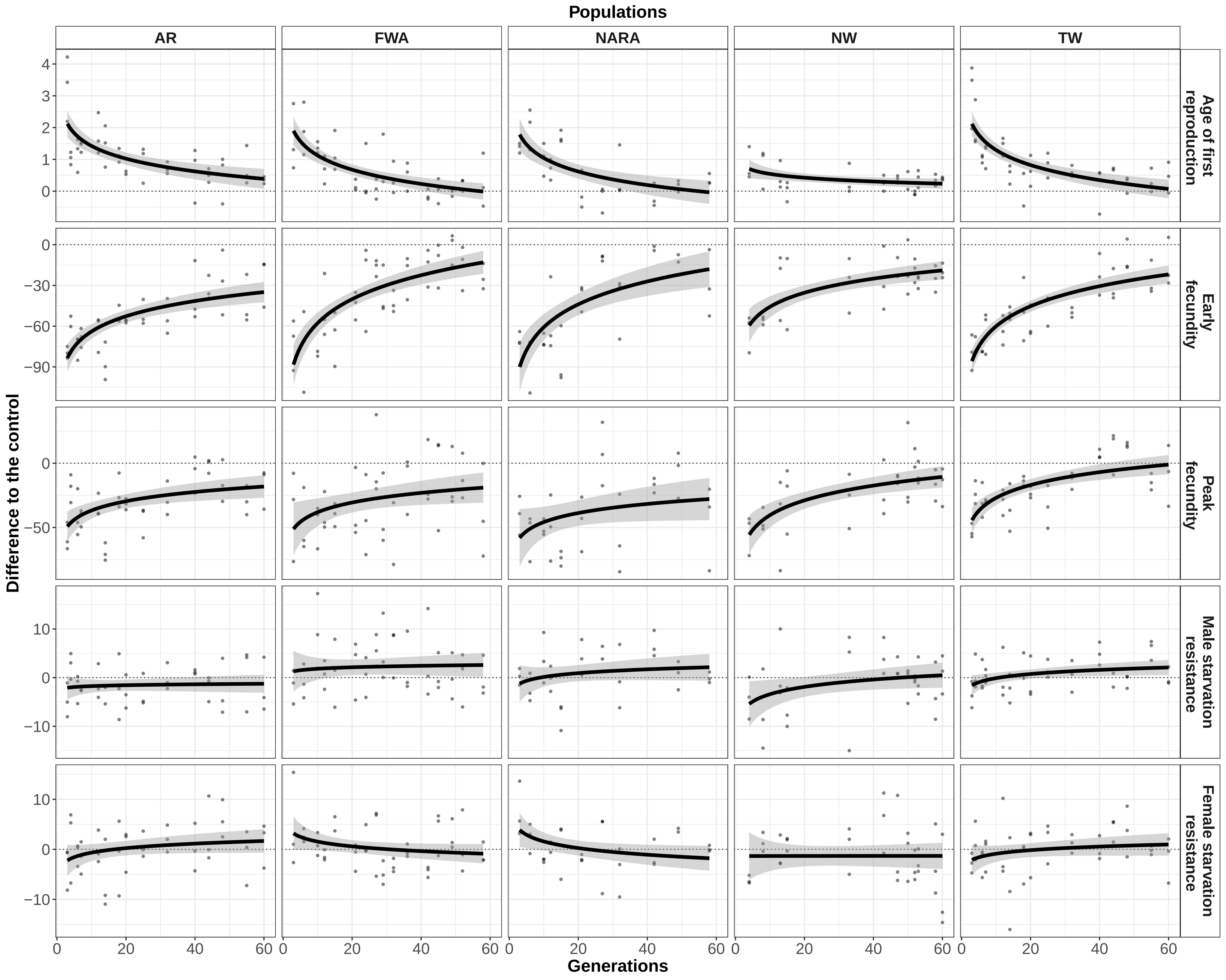
